# An Inducible T7 Polymerase System for High-Level Protein Expression in Diverse Gram-Negative Bacteria

**DOI:** 10.1101/2022.10.27.514131

**Authors:** Jennifer L. Greenwich, Melene A. Alakavuklar, Clay Fuqua

**Affiliations:** Department of Biology, Indiana University, Bloomington, IN 47405

**Keywords:** protein expression, T7 polymerase

## Abstract

A broad host range (BHR) inducible T7 RNA Polymerase system was developed, enabling induction with IPTG, similar to the *Escherichia coli* strain BL21(DE3), but now applicable in a wide range of bacteria. This allows for high protein yields and purification from diverse gram-negative bacteria including the native host.

---

High-level expression of proteins for purification is commonly performed using *Escherichia coli* BL21 (DE3). In addition to its native RNA polymerase, this strain harbors a prophage (DE3) that contains gene *1*, encoding the T7 RNA polymerase. The T7 polymerase recognizes a specific promoter sequence (*P*_T7_, TAATACGACTCACTATAG), and its own expression is under the control of the inducible *lacUV5* promoter, (1)). Similarly, ectopic expression of proteins in their native hosts is often used to study the role and effect of proteins, with expression of the protein being under control of the *lac* promoter. While these systems are generally sufficient, there are multiple drawbacks to each. For example, purification of protein from the native host is often limited, and the *lac* promoter (*P*_*lac*_) often does not provide sufficient expression. Purification of heterologous proteins from *E. coli* intrinsically removes them from their native physiological context, and hence any endogenous modification or regulation. Here we develop a bipartite plasmid-based *P*_T7_ expression system using the T7 polymerase. Similar to standard *E. coli* protein expression protocols, the gene of interest is expressed from the T7 promoter, in this case carried on a pVS-based (spectinomycin resistance, Sp^R^) BHR plasmid (2). The expression plasmid also provides an efficient N-terminal secretion signal derived from the *E. coli dsbA* gene (3), that can be used for secretion of the expressed proteins. A second compatible plasmid (pBBR origin, gentamycin resistance, Gm^R^) provides the T7 polymerase gene, expressed from *P*_*lac*_ and carrying *lacI*^*Q*^, and thus effectively regulated with IPTG (4, 5). An alternate T7 polymerase expression plasmid was also created with an IncP origin and a tetracycline resistance (Tc^R^) marker (6) and is also available.

The inducible T7 polymerase plasmid, pJLG038, is a BHR derivative of pSRKGm (a derivative of pBBR1MCS), (4, 5) and carries *gnt*, conferring gentamicin resistance. To construct the plasmid, the *1* gene was amplified from BL21(DE3) and ligated into pSRKGm. The gene was amplified using the following primers (restriction enzyme cut sites underlined): forward-gtac<u>tctaga</u>atgaacacgattaacatcgc; reverse-gtac<u>gtcgac</u>ttacgcgaacgcgaagtccg and ligated into pSRKGm as a *XbaI* and *SalI* fragment. The IncP plasmid, pJLG039, was constructed in a similar manner. From there, the plasmid was purified and sequenced before being electroporated into *A. tumefaciens* for protein induction experiments. The three plasmids can also be introduced by conjugation through an IncP delivery system. See Figure 1A-C for gene maps.

**Figure 1:**
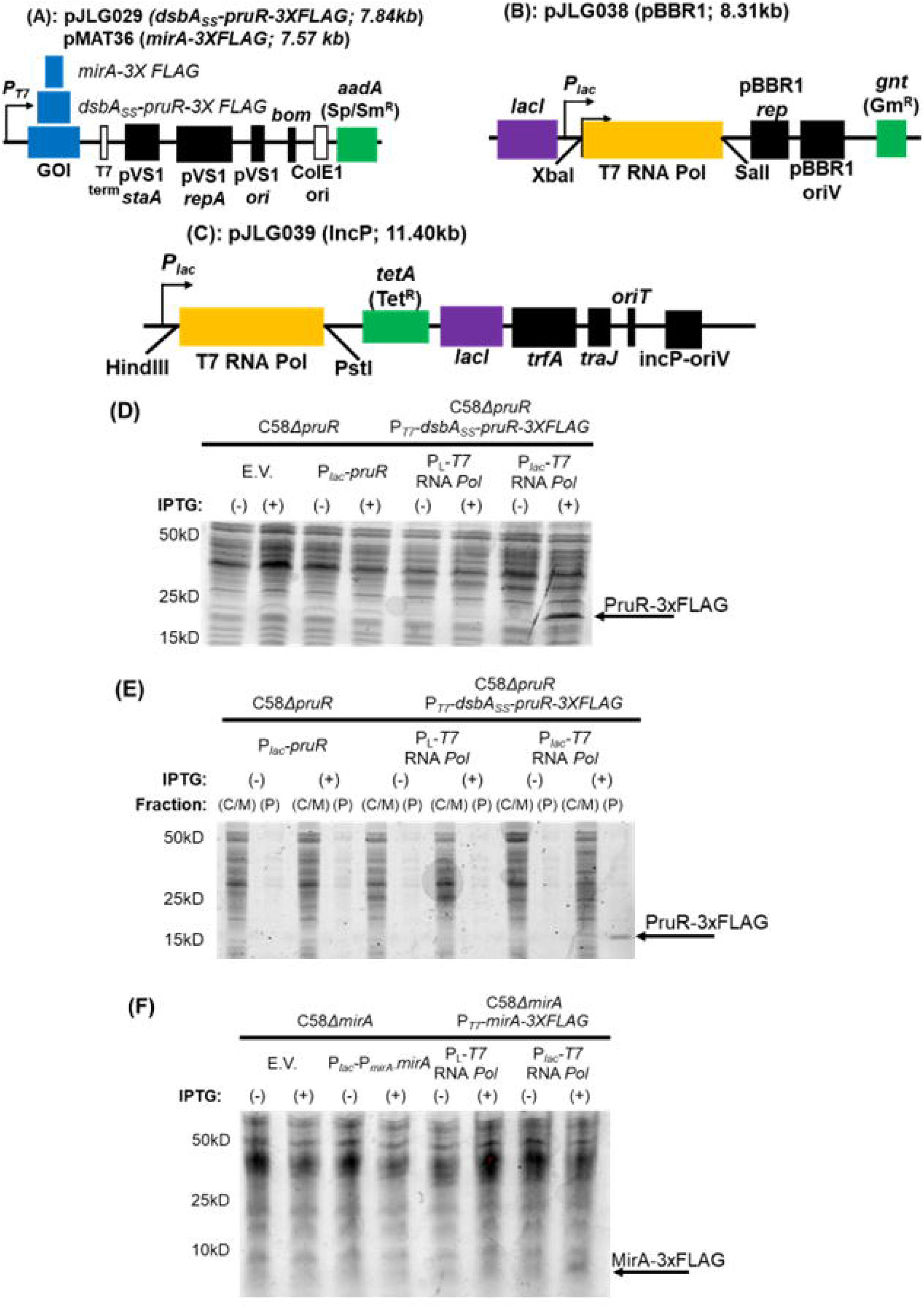
Plasmid Maps and Protein Production in *A. tumefaciens*. Boxes with text indicate relative position of genes. (A): Map of pJLG209 and pMAT36. The gene of interest is indicated in blue and the T7 terminator sequence in white. pVS1 *staA*, pVS1 *rep*, and pVS1 *ori* are a plasmid stability gene from the pVS1 plasmid, a replication protein, and the origin of replication, respectively (black). *Bom* is the basis of mobility gene and necessary for conjugation. The ColE1 ori is active in *E. coli* (white), and *aadA* encodes an aminoglycoside transferase (green), conferring resistance to spectinomycin and streptomycin. (B-C): Maps of pJLG038 and pJLG039. *lacI* (purple) encodes the lactose repressor, T7 polymerase (gold) is ligated at the indicated sites; The origin, replication proteins, and conjugation proteins are indicated by black boxes; pBBR1 rep and pBBR1 ori encode the replication protein and origin, for pJLG038, *traJ* (for conjugation) and *incP-oriV*, and *oriT* for pJLG039. Antibiotic resistance genes (green): *gnt* encodes gentamicin acetyltransferase and *tetA* encodes a tetracycline efflux pump. (D): Expression of the periplasmic protein PruR using either an untagged version expressed by the *lac* promoter or a 3XFLAG tagged version of the protein expressed by the T7 promoter. PruR is detectable in the strain with the IPTG-inducible *lac* promoter driven T7 RNA polymerase. (E): Periplasmic fractionation of the strains used in A (except for the negative control) shows that only in the inducible T7 RNA polymerase strain is there sufficient protein to be visible by SDS-PAGE. (F): Expression of MirA, a cytoplasmic protein, using either the *lac* promoter fused to its native promoter (due to low protein expression from just the *lac* promoter) and a 3XFLAG tagged version of the protein under the control of the T7 promoter. As with PruR, noticeably more protein is seen with the inducible T7 RNA polymerase. E.V., empty vector; P_L_-T7 polymerase, constitutively expressed T7 polymerase; (C/M), cytoplasmic/membrane fraction; (P), periplasmic fraction.

In addition to the inducible T7 plasmid, a second compatible plasmid with a different antibiotic resistance marker carrying the protein of interest is required, often with an affinity tag to aid subsequent purification. Here we used derivatives of pRA301 (2) constructed using isothermal assembly. The protein of interest can be fused to a 3XFLAG tag for protein purification and which may also be useful to determine protein levels using anti-FLAG antibodies.

While this work was performed and validated in *Agrobacterium tumefaciens*, the plasmids contain BHR origins of replication and are compatible with a diversity of Gram-negative taxa. With the aim to purify protein directly from *A. tumefaciens*, we discovered that available expression systems (*P*_*lac*_, *P*_N25_, *P*_*traI*_) were insufficient for high-level protein expression (4, 7, 8). An existing system provided constitutive expression of the T7 polymerase (9), enabling constitutive protein expression in *A. tumefaciens*. However, we observed surprisingly weak expression and we hypothesized that the constitutively expressed T7 polymerase negatively impacts cellular physiology, and selects for mutants with decreased target protein expression. There is evidence of the T7 polymerase gene accumulating mutations in BL21(DE3), leading to lower protein production levels (10).

In contrast to these systems, our IPTG-inducible T7 polymerase expression system, produced high levels of protein that was strictly dependent on IPTG (Fig. 1D). No growth inhibition was observed during induction for the proteins we tested. We initially designed and implemented this system to purify the periplasmic PruR protein (11), and periplasmic fractionation confirms that the protein is efficiently targeted to the periplasm, and at higher levels than with other protein expression systems (Fig. 1E). The system is also effective for expression of the MirA cytoplasmic protein as well (Fig. 1F) (12).

## Data availability

The complete sequences of plasmids pMAT36, pJLG029, pJLG038, and pJLG039 have been deposited into GenBank (accession numbers <u>OP627879</u>, <u>OP627880</u>, <u>OP627881</u>, <u>OP627882</u>) and are available from Addgene (ID numbers 19279, 12980, 12981, 12982).

## Acknowledgements

This project was supported by National Institutes of Health (NIH) grant GM120337 (CF).

